# Loss Of function of Male-Specific Lethal 3 (Msl3) Does Not Affect Spermatogenesis in Rodents

**DOI:** 10.1101/2023.03.16.532933

**Authors:** T.A. Mitchell, J.M. Lin, S.M. Hicks, J.R. James, P. Rangan, P.E. Forni

**Affiliations:** Department of Biological Sciences, University at Albany, State University of New York, Albany, NY 12222, USA; The RNA Institute, University at Albany, State University of New York, Albany, NY 12222, USA; The Center for Neuroscience Research, University at Albany, State University of New York, Albany, NY 12222, USA; Black Family Stem Cell Institute, Department of Cell, Developmental, and Regenerative Biology, Icahn School of Medicine at Mount Sinai, 1 Gustave L. Levy Place, New York, NY 10029, USA

## Abstract

Msl3 is a member of the chromatin-associated male-specific lethal MSL complex which is responsible for the transcriptional upregulation of genes on the X chromosome in males Drosophila. Although the dosage complex operates differently in mammals, the Msl3 gene is conserved from flies to humans.

Msl3 is required for meiotic entry during Drosophila oogenesis. Recent reports indicate that also in primates, Msl3 is expressed in undifferentiated germline cells before meiotic entry. However, if Msl3 plays a role in the meiotic entry of mammals has yet to be explored. To study this, we used mouse spermatogenesis as a study model. Analyses of single cells RNA-seq data revealed that, in mice, Msl3 is mostly expressed in meiotic cells.

To test the role of Msl3 in meiosis, we used a male germline-specific Stra8-iCre driver and a newly generated Msl3^flox^ conditional knock-out mouse line. Msl3 conditional loss-of-function in spermatogonia did not cause spermatogenesis defects or changes in the expression of genes related to meiosis. Our data suggest that, in mice, Msl3 exhibits delayed expression compared to Drosophila and primates, and loss-of-function mutations disrupting the chromodomain of Msl3 alone do not impede meiotic entry in rodents.

## Introduction

Gametes ensure the survival and propagation of all sexually reproducing organisms. Gametes arise from germ cells that differentiate and undergo meiosis to produce haploid oocytes and sperms. Meiosis is essential for reductional cell division to generate haploid gametes and genetic diversity by promoting recombination. Thus, regulation of meiosis is critical for gametogenesis and fertility. However, the regulation of meiotic entry remains incompletely understood.

Mammalian spermatogenesis is an excellent model system to determine regulators of meiotic entry due to the presence of spermatogonial stem cells (SSCs) that self-renew and differentiate, undergoing meiosis throughout the animal’s life [1, 2]. This differs from oogenesis, where most of the germ cells directly differentiate into oocytes during embryonic development that then meiotically arrests [3, 4].

Spermatogenesis in mammals initiates with SSC, which are also called Single spermatogonia (A_s_), which give rise to A paired spermatogonia (A_paired_) and A aligned spermatogonia (A_aligned_), which make chains of 4, 8 and 16 Type A spermatogonia which eventually gives rise to type B differentiated spermatogonia through mitotic divisions [5]. Type B spermatogonia enter meiosis prophase I when they initiate recombination. These type B spermatogonia then progress to a second round of meiotic divisions to yield round spermatids. Round spermatids then go through spermiogenesis, in which morphological changes occur, including the trading of histones for protamines, to get elongated spermatids and eventually mature spermatozoa [5-7].

In mammals, the entry into meiosis is promoted by steroid signaling during oogenesis and spermatogenesis. Retinoic acid (RA) from somatic cells surrounding the spermatogonia activates the transcription factor *Stimulated by retinoic acid 8* (*Stra8*) in the germline [8-13]. Stra8 fosters meiotic entry during spermatogenesis by promoting the transcription of a broad gene expression program [14, 15]. However, *Stra8* is not sufficient to induce meiosis, suggesting a cell type-specific chromatin landscape and/or factors that cooperate with *Stra8* [16].

Male-specific lethal 3 (Msl3) is a member of the Male-specific lethal (MSL) complex which is necessary for dosage compensation in *Drosophila [17]*. Dosage compensation promotes transcriptional upregulation of genes on the X chromosome in males to equal the two copies from the two X chromosomes in females [18, 19]. This transcriptional upregulation is essential to the survival of male *Drosophila*. The compensation complex comprises several components, including MSL1, MSL2, Msl3, male-absent on first (MOF), maleless (MLE), and two roX noncoding RNAs [20]. Msl3’s chromodomain is essential for binding to X-linked genes by recognizing Histone 3 lysine 36 trimethylation marks (H3K36me3) and H4K20 monomethyl marks [17, 21-23].

During *Drosophila* oogenesis Msl3 is expressed from the undifferentiated germline stem cells (GSC) to the differentiating cysts stages when the germ cells enter meiosis [24]. In *Drosophila* Msl3 functions as part of a transcriptional axis that promotes GSC differentiation and meiotic entry by recognizing H3K36me3 marks, presumably via its chromodomain [24].

A recent research report [25] has identified Msl3 expression in undifferentiated spermatogonia of primates and humans, suggesting its potential role in the germline in mammals. Furthermore, large-scale analyses of the X chromosome in infertile men have identified Msl3 as one of the genes associated with spermatogenic failure [25-27]. These findings provide evidence that Msl3 may indeed play a significant role in meiotic entry in mammals.

Through the analysis of published single-cell (sc) RNA-sequencing data [28], we have discovered that Msl3 expression mostly occurs in meiotic cells, which differs from the previously reported expression pattern in primates and humans, where Msl3 was described in undifferentiated cells prior to meiotic entry [25, 27]. To further investigate the role of Msl3, we generated a new Msl3 conditional knockout mouse model. Interestingly, we found that disruption of the chromodomain of Msl3 [17] does not disrupt spermatogenesis in rodents. These mutant mice exhibit normal viability and fertility, suggesting that the chromodomain of Msl3 may not be essential for spermatogenesis in rodents.

## Materials & Methods

### Animals

Mice used in this study were as follows: Msl3 floxed (Msl3^em1Forni^), Stra8-iCre [29] and R26tdTomato [30]. All mice described were maintained on a C57BL/6J background. The Msl3 floxed mouse line (Msl3^em1Forni^) was designed using CRISPR/Cas9 technology and generated for us by Cyagen. This mouse line has been recently donated to The Jackson Laboratory. The Msl3 gene consists of 13 exons, and exons 2-4 was targeted using CRISPR/Cas9 inserting Lox*P sites* flanking exons 2 and 4 (Fig.3). The targeting vector was engineered with homology arms and the conditional knock-out region generated by BAC clone RP24-103L17 from the C57BL/6N library as a template. Cas9 mRNA was injected into fertilized eggs along with Msl3 gRNA and the targeting vector. Founder mice were identified by PCR and bred to C57BL/6N to test germline transmission. The Stra8-iCre (stock #017490) [29], and R26tdTomato (stock #007914) [30] mouse lines are commercially available from JAX.

Genotyping of the mouse lines was conducted by PCR using tail genomic DNA. iCre Primers: *Fwd*-AGA TGC CAG GAC ATC AGG AAC CTG, *Rev*-ATC AGC CAC ACC AGA CAC AGA GAT, *Fwd control-*CTA GGC CAC AGA ATT GAA AGA TCT, *Rev control*-GTA GGT GGA AAT TCT AGC ATC ATC C. Msl3^flx^ Primers: *Fwd-*TTCATCGGTCTAGGTGTAACTCCA, *Rev-* GAAGGCTCTGCTACAGTCTGATAC, for Msl3^flx^ targeted allele after Cre recombination we used Fwd-TTCATCGGTCTAGGTGTAACTCCA, *Rev-* GAAGGCTCTGCTACAGTCTGATAC. R26tdTomato Primers: *IMR0920-*AAGGGAGCTGCAGTGGAGTA, *IMR09021-* CCGAAAATCTGTGGGAAGTC, *IMR9103*-GGCATTAAAGCAGCGTATCC, *IMR9105*-CTGTTCCTGTACGGCATGG.

All animal experiments were conducted in accordance with the guidelines of the Institutional Animal Care and Use Committee (IACUC) at the University at Albany, SUNY.

### Surface spreading and Immunofluorescence

Testes were excised from male mice euthanized at P17-20. Excised testes were placed in ice-cold PBS. Testes were then weighed to determine incubation time in hypotonic extraction buffer (HEB). 30 mg or less for 30 min, tubules from a pair of testes weighing 31 to 50 mg for 30 - 45 minutes, and larger testes for 60 minutes. Testes were then placed back in ice cold PBS to remove the tunica albuginea and extract the seminiferous tubules. Seminiferous tubules were placed in HEB consisting of 30 mM Tris pH 8.2, 50 2 mM Sucrose, 17 mM Trisodium Citrate Dihydrate, 5 mM EDTA and incubated according to their weight. Seminiferous tubules were then transferred back into a petri dish and separated into small clumps. Individual clumps were placed in 60μl droplets of 100 mM sucrose pH 8.2 and repeatedly pipetted until the solution became turbid. Droplets of seminiferous tubules were placed in the corner of uncharged slides previously soaked in a fixative solution of 1% PFA and 0.15% triton X-100 pH 9.2 with a droplet of fixative remaining on the slide [31, 32]. The droplet was allowed to spread along the slide then left to dry overnight in a humidified chamber at 4°C. Slides were then stored at -80°C until use.

For immunofluorescence analysis, the spreading nuclei were immunostained with mouse anti-SYCP3 (1:500), rabbit anti-SYCP1 (1:500) and mouse anti-γH2AX (1:500). Using SYCP3 to label axial/lateral element characteristics and DAPI-stained heterochromatin pattern, spermatocytes in spreading were classified as leptotene, zygotene, pachytene and diplotene.

### Histological and immunohistochemical analysis

Testes were fixed in 4% formaldehyde embedded in OCT and sectioned, then stored at -80°C until use. For immunofluorescence analysis, the following primary antibodies were used: Rabbit anti-RFP (1:500 Rockland), Rabbit anti-Stra8 (1:250 ABCAM), mouse anti-SCP3 (1:500 ABCAM).

Testes weight and Seminiferous Tubule Diameter Measurements. Average weights were calculated by weighing testes (n=3 for each genotype) immediately after excision and averaged for each genotype. Average seminiferous tubule diameter measurements were calculated by measuring 10 seminiferous tubules per animal per genotype and calculating the average diameter of the 10 seminiferous tubules. This was repeated for a total of 3 animals per genotype. The average of the 3 averages was taken for each genotype as the representative weight.

### RNA-sequencing

For whole RNA sequencing, testes were collected, and RNA extraction was performed using the Thermo Fisher RNA PureLink Mini Kit. Sequencing was performed using Illumina Nextseq 500.

For Single-cell RNA sequencing, we utilized previously published data by Green at al. [28] available through NCBI’s Gene Expression Omnibus, GEO series accession number GSE112393 (https://www.ncbi.nlm.nih.gov/geo/query/acc.cgi?acc=GSE112393). Files were imported into the R package *Seurat 4*.*0*.*5 [33]* for filtering. After filtering, mitochondrial genes and RNA features, individual files were merged into one file representing all germ cell types. Dimensionality reduction was performed using principal component analysis (PCA).

### Gene Ontology Term Analysis

Gene Ontology term analysis was performed using the Panther Classification System.

## Results

### Msl3 is expressed in mouse testes during meiotic stages via ScSeq

To determine when Msl3 is expressed in mouse testes, we turned to single-cell RNA-sequencing (scRNA-seq) analysis. Using a publicly available drop-seq data set (GSE112393 [28]) we reproduced a PCA clustering of testes germ cells. Markers used to identify spermatogonia were as follows: Zbtb16, Uchl1, Stra8, and Kit [10, 34, 35]. Markers used to identify spermatocytes were as follows: Sycp3, Tex101, Spag6 and H2afx [36-39]. Markers used to identify round/elongating spermatids were as follows: Acrv1, Tssk1, Tnp1, Prm1 [40-43] (Fig. 2A,B). As continuous transitions occur between cell states during spermatogenesis, they are represented by the changes in expression patterns of marker genes. (Fig. 1B).

**Figure 1.**
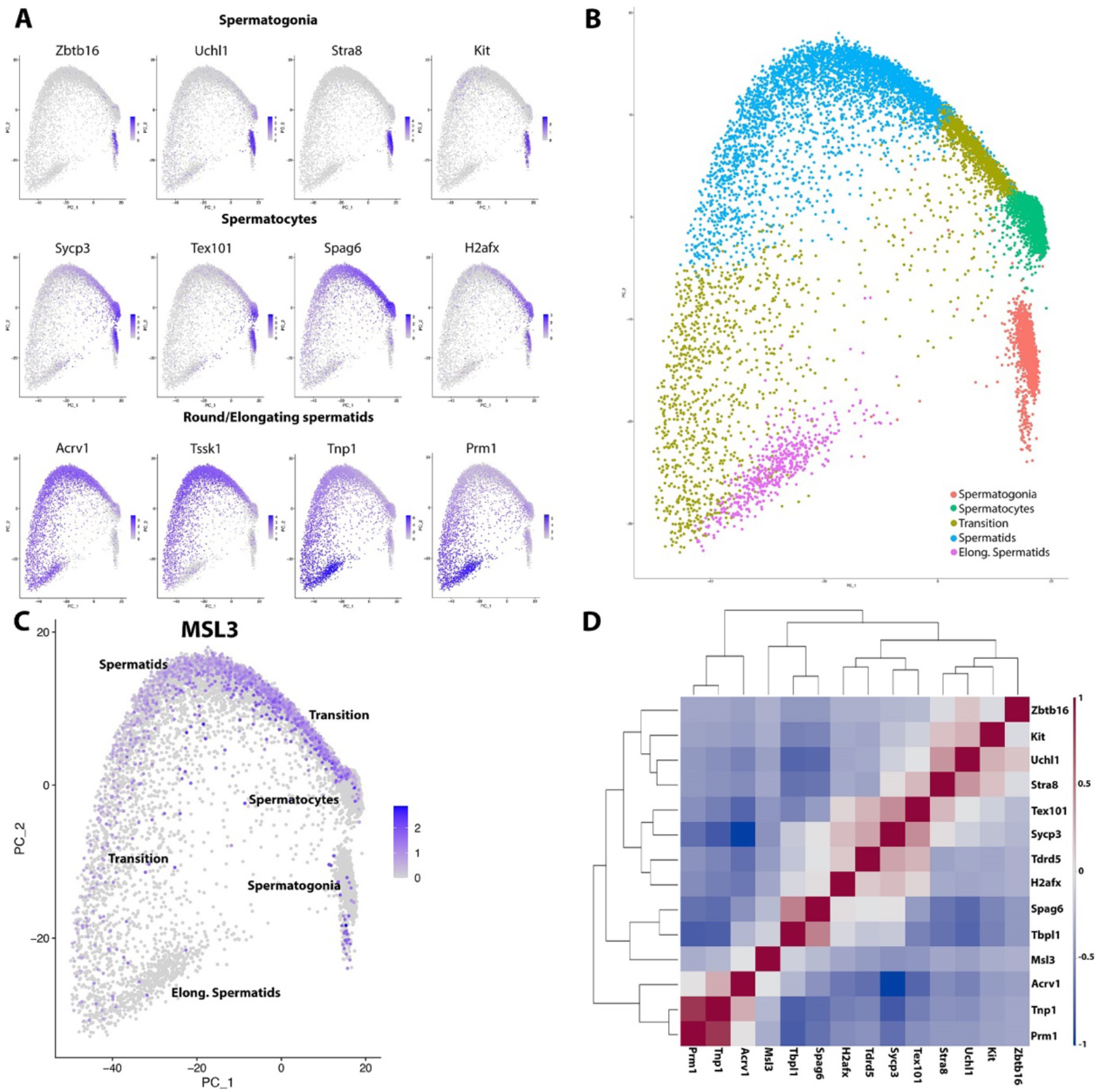
Single-cell RNA sequencing of mouse testes. A) PCA plot of different germ cell population markers for spermatogonia, spermatocytes and round/elongating spermatids to identify germ cell type of each cluster. B) Clusters grouped into spermatogonia, transition states, spermatocytes and spermatids based on the marker genes they expressed to establish the trajectory of spermatogenesis. C) Msl3 is expressed in spermatocytes and continues into spermatids then ceases as spermatids transition into elongating spermatids. D) Correlation map of different marker genes showing that Msl3 expression most closely correlates with that of Tbpl1 and Spag6 which are markers of spermatocytes.

**Figure 2.**
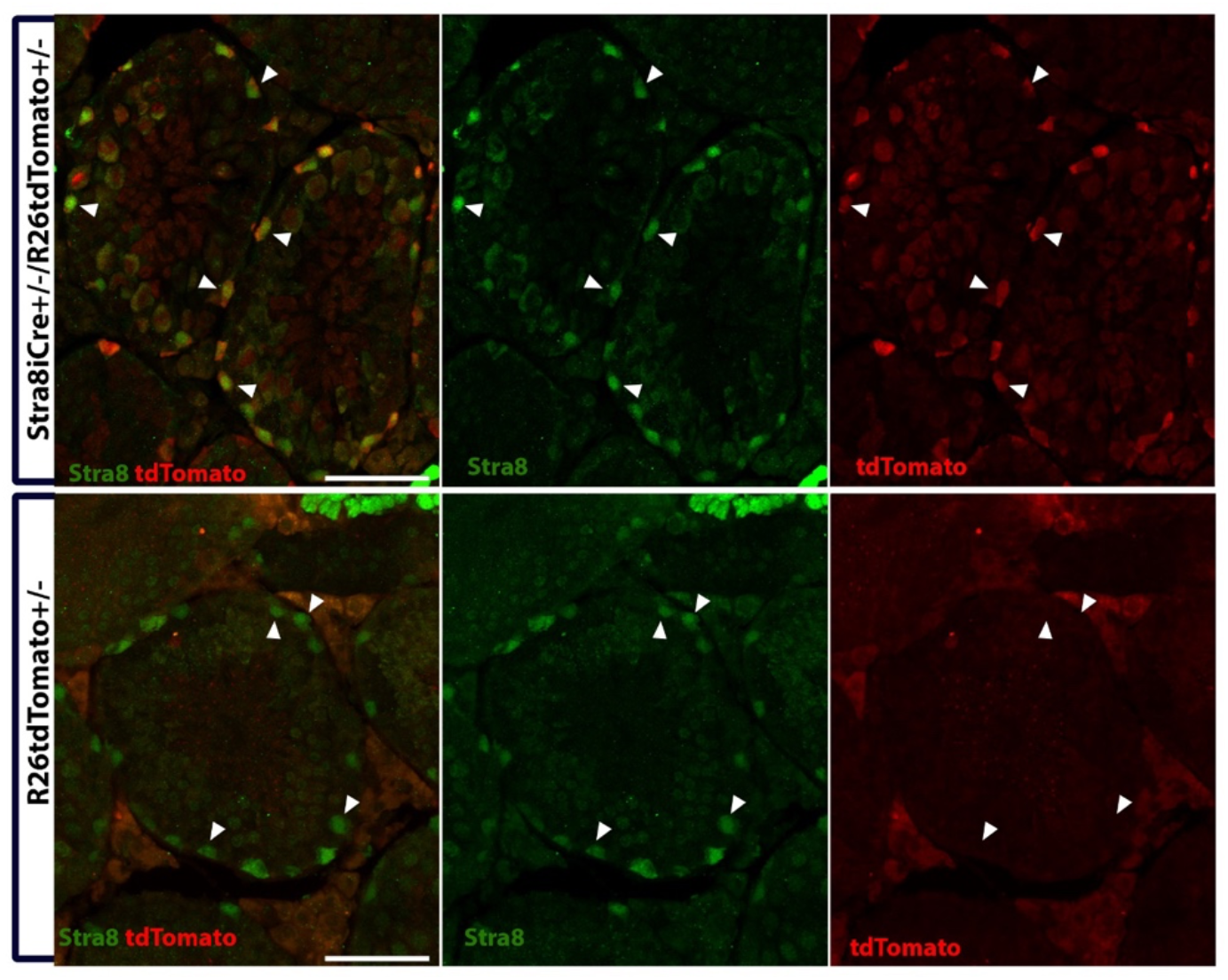
Immunofluorescence staining against Stra8 and tdTomato at p30. Stra8 in green, tdTomato in red. Immunofluorescence shows that in Stra8iCre^+/-^R26tdTomato^+/-^ mutants, Stra8 and tdTomato staining colocalize, while in tdTomato^+/-^ controls, Stra8 positive cells are negative for tdTomato. This confirms that tdTomato is only expressed in Stra8iCre cells. Scale bar = 50μm.

**Figure 3.**
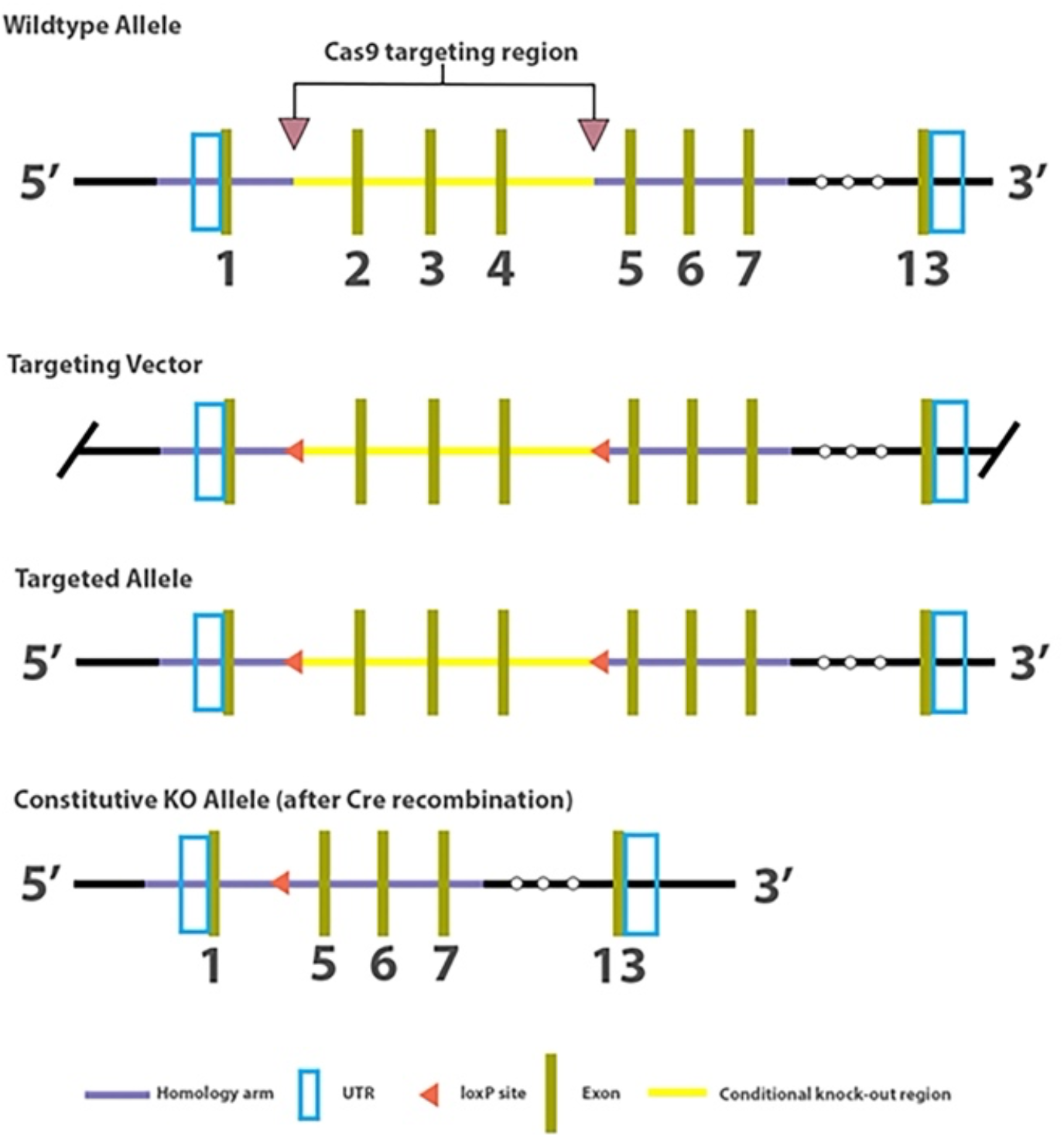
Strategy adopted for Msl3 conditional knock-out mouse line generation. Crispr/Cas9 was used to insert loxP sites upstream of exon 2 and downstream of exon 4 of the Msl3 chromodomain. Upon Cre recombinase activity, exons 2-4 are excised from the Msl3 gene generating a constitutive knockout allele.

After identifying germ cell populations, we analyzed the expression of Msl3. Our observations revealed limited Msl3 expression in spermatogonia (fig.1C). However, we noticed a consistent expression of Msl3 expression spanning from spermatocytes to round spermatids (fig.1C). This finding suggests that Msl3 is primarily expressed during the meiotic stages of mouse spermatogenesis. To further validate the expression pattern of Msl3, we examined its correlation with spermatocyte markers Tbpl1 and Spag6 (fig.1D) [44]. Our analysis confirmed Msl3 expression during mouse spermatogenesis (fig.1D).

### iCre successfully recombines in Stra8+ cells

To assess the efficacy of iCre recombination in the testes, we performed a cross between female Stra8iCre mice [29] and male R26tdTomato reporter mice. We examined the presence of tdTomato expression in Stra8-positive cells. Testes were collected from P30 R26tdTomato/Stra8iCre males and processed for fixation, sectioning, and immunostaining against tdTomato and Stra8. Immunostaining analysis revealed the colocalization of Stra8+ spermatogonia and tdTomato+ cells in Stra8iCre/R26tdTomato mice, while R26tdTomato control mice did not show such colocalization (refer to Figure 2). This observation confirmed the specificity of tdTomato expression in spermatogonia and demonstrated the successful recombination of iCre in Stra8+ cells. However, only low levels of tdTomato expression were detectable in the meiotic cells. By quantifying the Stra8+ and tdTomato+ cells, we determined a high recombination efficacy of 99.4% in the testes.

### Msl3 mutant mice show normal testes morphology

The mouse Msl3 chromodomain is encoded in exons 1-5 of the Msl3 gene on Chromosome X. The chromodomain has been proposed to play a key role in controlling the formation of the synaptonemal complex in drosophila [24]. To investigate the role of Msl3 in mouse spermatogenesis, we employed a conditional knockout strategy targeting exons 2-4 of the Msl3 gene.

Initially, we examined whether Msl3cKO (Stra8iCre+/-; Msl3flox/flox) mice exhibited any apparent testicular morphological defects. At P45, when males reach sexual maturity and all cell types are expected to be present in the testes, we dissected the testes and assessed their size, shape, and weight. We found no significant differences in these parameters between the control group (0.056 g +/-0.003) and the mutant group (0.059 g +/-0.002), indicating that gross testicular morphology was unaffected in Msl3cKO mice (Fig. 4A, B).

**A.) Figure 4.**
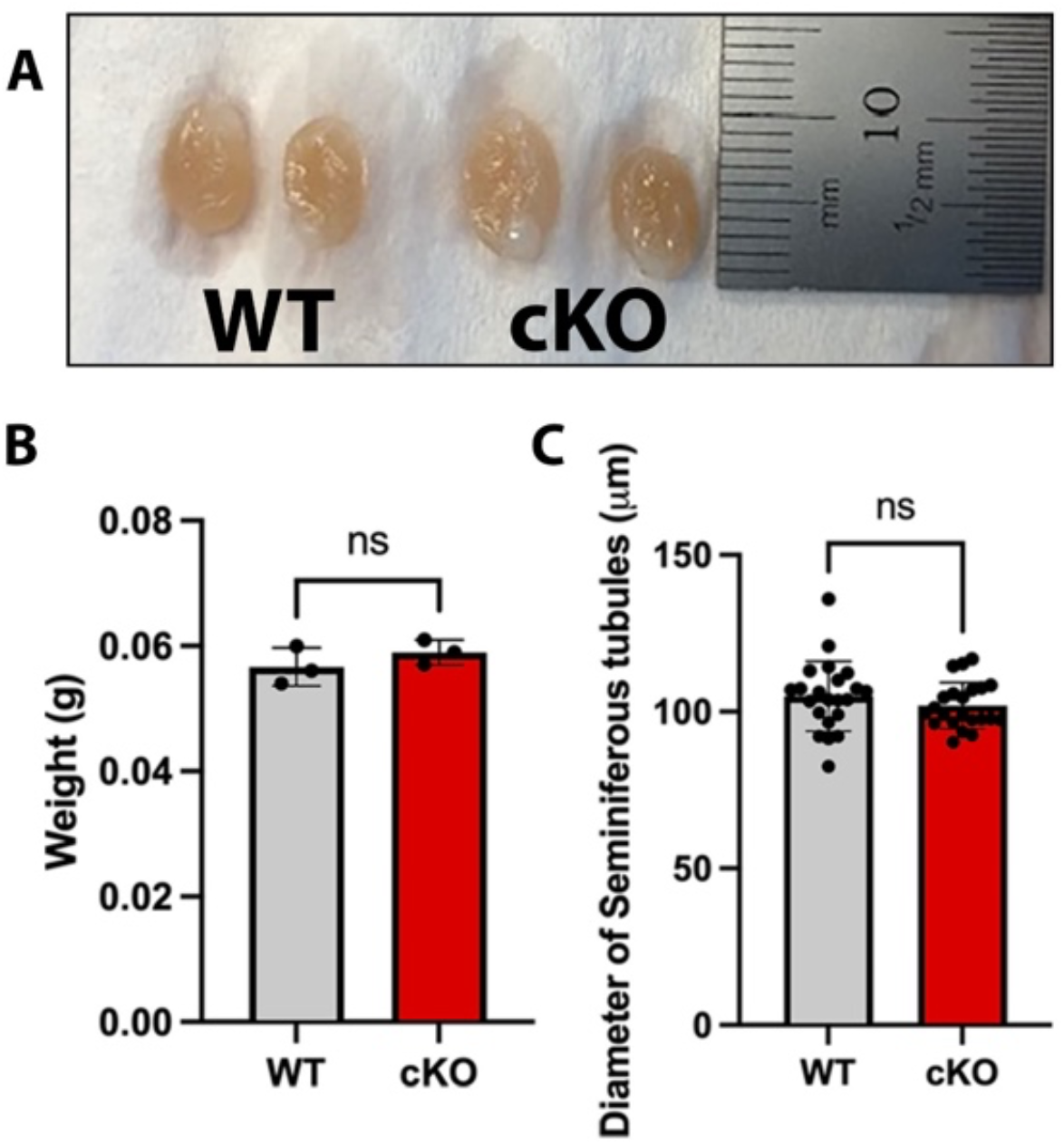
Image of Msl3 wildtype (WT) (left) and Msl3 cKO (right) testes at P45. Image shows comparable gross morphology, size, and shape. B.) Bar graph illustrating the weight of pairs of testes for Msl3 WT and Msl3 cKO mice. C.) Bar graph showing the average diameter of the seminiferous tubules for Msl3 WT and Msl3 cKO mice (10 seminiferous tubules per animal, n=3 animals per genotype).

Further examination of testes sections under a microscope revealed no morphological defects in either the mutants or controls. Additionally, measurements of seminiferous tubule diameter showed no significant differences between the two groups (controls: 114.6 μm +/-4.6; mutants: 115.0 μm +/-6.7) (Fig. 4C). Both mutant and control testes displayed rounded seminiferous tubules with comparable cellular organization, and all cell types, ranging from primary spermatogonia to elongated spermatids, were observed based on the morphology of cells within the seminiferous tubules.

### Formation of synaptonemal complex in the absence of functional Msl3

The scRNA-Seq analysis revealed the expression of Msl3 in primary spermatocytes (Fig. 1). Previous studies have suggested Msl3 involvement in the transcriptional upregulation of synaptonemal complex proteins [24]. To investigate whether Msl3 is necessary for meiosis in mice, we examined the meiotic prophase-I stage using a marker for synapsis facilitated by synaptonemal complex proteins.

Meiotic chromosome spreads were prepared from P17-21 control and Msl3 cKO mutant mice. Immunofluorescent staining was performed using an antibody against SYCP3, one of the lateral elements of the synaptonemal complex [45]. Based on the staining pattern, primary spermatocytes were characterized as being in leptonema, zygonema, pachynema, or diplonema stages. Both control and Msl3 cKO chromosome spreads showed spermatocytes at all stages of prophase I, indicating that both genotypes could complete prophase I.

Quantitative analysis of spermatocytes from both mutants and controls revealed no significant differences in the distribution of spermatocytes between the two genotypes (Fig. 5). This analysis was based on changes in SYCP3 localization [46] transitioning from a more diffuse or widespread expression in leptonema to a condensed expression with strong puncta at the centromeric region in diplonema.

**Figure 5.**
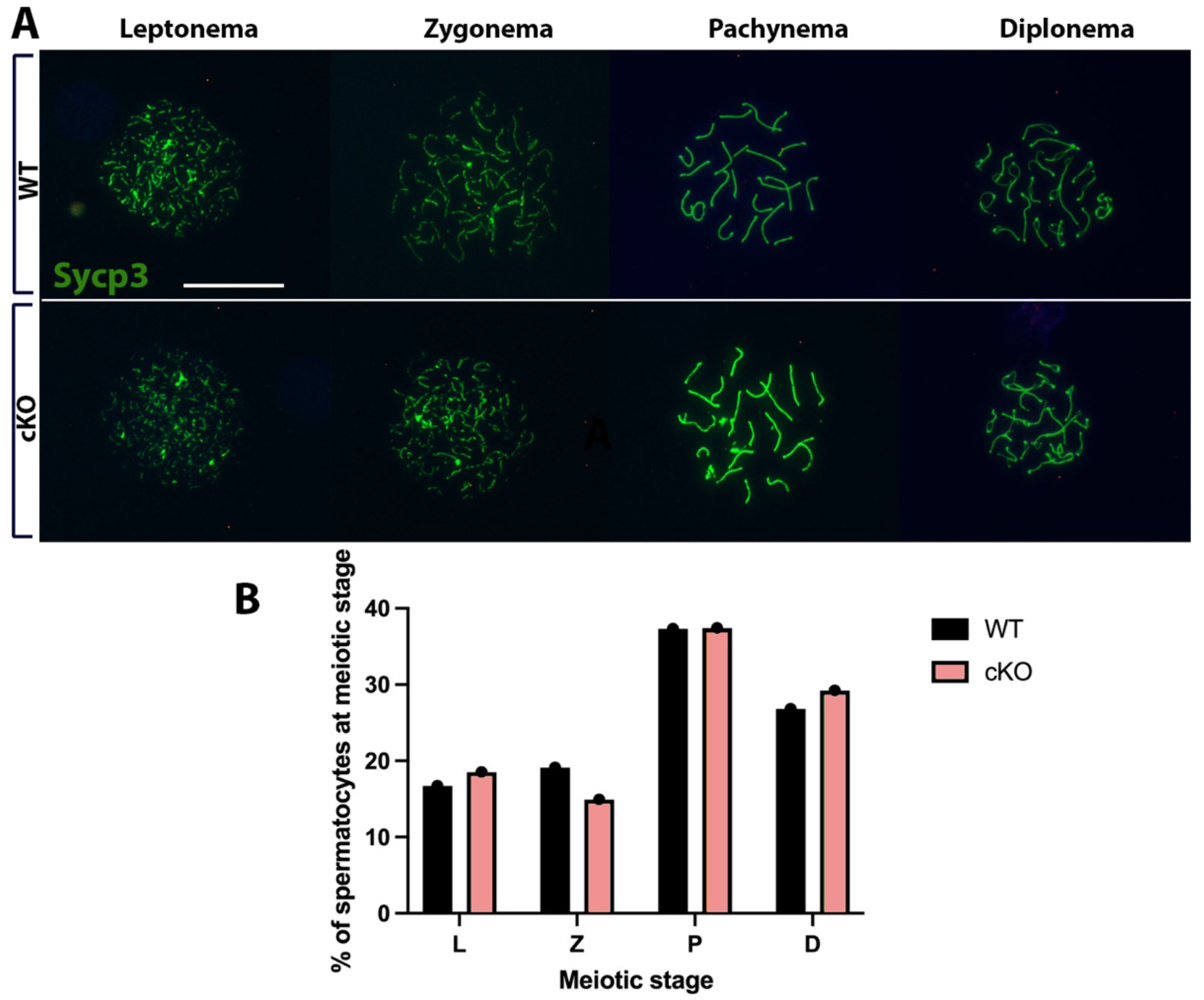
A.) Meiotic chromosome spread of spermatocytes from P17-21 Msl3 WT and cKO testes stained with antibody against SYCP3. Meiosis prophase I was examined between mutants and controls. Spermatocytes were characterized as in leptonema, zygonema, pachynema or diplonema. Scale bar=25um B.) 209 spermatocytes were counted for WT controls and 195 for Msl3 cKO. Subphases of meiosis prophase I show no significant differences between Msl3 WT and Msl3 cKO. N=3 per genotype

These findings suggest that, in rodents, differently from Drosophila, Msl3 is not necessary to form the synaptonemal complex, [24].

### Msl3 mutants are fertile and produce normal litter sizes

Histological analysis of Msl3 cKO seminiferous tubules showed the presence of all cell types, from primary spermatogonia to elongated spermatids. The presence of elongated spermatids showed that cells could initiate spermiogenesis leading to the formation of mature spermatozoa. Msl3 cKO males were thus mated with WT females to examine their fertility along with littermate controls. Each Msl3 cKO male was paired with 2 C57B/6J females and allowed to mate for 2 weeks. Msl3 cKO males successfully impregnated both females and produced litters of comparable sizes to that of controls. Msl3 cKO offspring were also viable (Table 1).

**Table 1.** Msl3 cKO males and littermate controls were mated with 2 C57B/6 WT females each. Males were given 2 weeks to impregnate females and removed after the 2-week period. All males successfully impregnated respective female. The average litter size of Msl3 cKO males (8.17± 1) was not significantly different from that of littermate Msl3 WT controls (8.5± 1.05) (p=0.5826).

| Genotype | # of Breeders | # of Litters | Avg. litter size |
| --- | --- | --- | --- |
| MSL3 WT | 3 | 6 | 8.5 ± 1.05 |
| MSL3 cKO | 3 | 6 | 8.17 ± 1 |

### Loss of Msl3 shows no gross disruptions in the transcriptome

As our Msl3 mutant mice did not exhibit any defects in spermatogenesis, we investigated whether the upregulation of other genes could compensate for the potential function of Msl3. To address this, we performed whole RNA-sequencing on Msl3 cKO mutants and wildtype controls, with three replicates for each genotype. Differential gene expression (DGE) analysis was conducted on the sequenced testes to identify any genes that were differentially expressed between Msl3 cKOs and controls.

Visualization of read counts for Msl3 between genotypes using the Interactive Genomics Viewer showed a few reads mapping to exons 2-4 in our Msl3 cKOs (Fig. 6A), consistent with the deletion of most of the chromo domain. Notably, only a few read counts were observed in our cKOs, which could reflect Msl3 expression in Sertoli cells or in spermatogonia before Stra8-mediated recombination.

**Figure 6.**
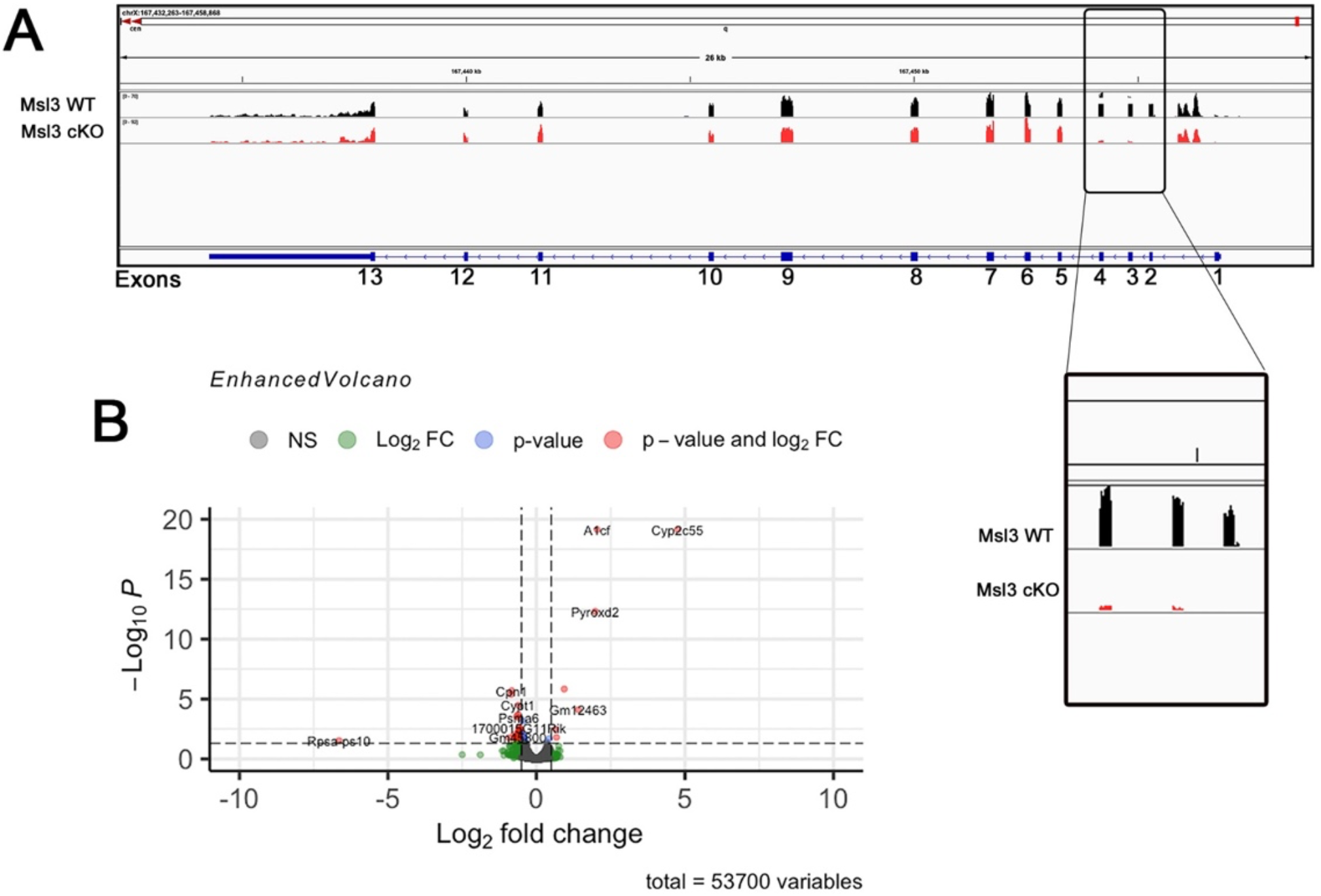
**A)** Read counts between Msl3 WT and Msl3 cKO show little to no reads for exons 2-4, validating their excision from the gene in our cKO model. **B.)** Volcano plot showing significantly upregulated and downregulated genes in our Msl3 cKO model.

The differential expression analysis between control and mutant samples revealed the upregulation of seven genes and the downregulation of 44 genes in Msl3 cKOs (Fig. 6B). Genes were considered significant based on a log2 fold change >0.5 or <-0.5 and a p-adj. value <0.05. Among the seven upregulated genes, three (Cyp2c55, A1CF, Pyroxd2) showed the highest log2 fold change (0.5 or greater). These genes are involved in enzymatic and RNA metabolic functions; however, their reported functions in the testes or spermatogenesis are not well-established.

Of the 44 significantly downregulated genes, 18 exhibited a log2 fold change of -0.5 or lower. Gene ontology (GO) term analysis of genes with a log2 fold change of -0.5 or lower and 0.5 or greater did not reveal any convergence on commonly known pathways. Therefore, based on our analysis, the absence of functional Msl3 does not appear to alter the meiotic program during mouse spermatogenesis.

## Discussion

Our efforts to uncover a functional role of Msl3 in murine spermatogenesis revealed its dispensability in this context. Notably, there are expression pattern differences between rodents and primates, and humans. In humans, Msl3 was shown to be a marker of spermatogonia stem cells (SSC), particularly State 0 SSCs, in fetal-infant testes [27]. Similarly, in macaque monkeys, Msl3 is expressed in Spermatogonia state 1 (SPG1) [25]. Our study suggests that, in mice, Msl3 expression is mostly restricted to the spermatocyte stage and only in a small portion of spermatogonia (Figure 1C). In this study, we disrupted the chromodomain of Msl3 downstream of Stra8, which marks the differentiation phase of spermatogonia; we cannot exclude that Msl3 may be potentially critical before the differentiation of spermatogonia.

While Msl3 is important for transcriptionally upregulating synaptonemal complex components during *Drosophila* oogenesis, it does not appear to affect synaptonemal complex formation during mouse spermatogenesis. Using our Msl3 conditional knock-out mouse model, we observed no noticeable defects or interruptions in spermatogenesis.

The Msl3 gene contains a chromodomain and an MRG domain [47]. The MRG gene family, consisting of MRG15 and MRGX, is involved in cellular proliferation and cycle progression [48]. In *Drosophila*, the MRG domain mediates Msl3’s incorporation into the Dosage Compensation Complex [49]. The chromatin-binding protein MORF-related gene on chromosome 15 (MRG15) is involved in pre-mRNA splicing at the round spermatid stage [50].

Our conditional knockout lacks a functional chromodomain. It is possible that MRG and chromodomains serve redundant functions resulting in the lack of any observable phenotype in our mouse models. The diverse involvement of MRG genes leaves open this possibility of a role of the MRG domain in Msl3 function. It may be responsible for chromatin binding and histone acetyltransferase (HAT) [49] activity in transcriptionally active genes. However, while this manuscript was in preparation, the International Mouse Phenotype Consortium (IMPC) published on the IMPC website the phenotypic characterization for a constitutive Msl3 KO mouse line (MMRRC Strain #042195-JAX). Differently from the mouse line we generated, this constitutive Msl3 KO carries a deletion of exon 5, which leads to the early termination 127 amino acids later and the complete loss of the MRG domain. Surprisingly, also this Mls3 KO line resulted in being viable and without fertility defects.

In conclusion, our study suggests that loss-of-function of the chromodomain in premeiotic cells does not prevent meiotic entry or progression in rodents.

The observed differences in Msl3 expression and the phenotypic outcomes following its loss-of-function between Drosophila [24] and mice highlight potential evolutionary differences that may have arisen in the mechanisms governing meiotic entry and progression across species. These findings indicate that the regulatory pathways involving Msl3 in these processes have likely diverged during evolution, leading to species-specific variations in the roles and expression patterns of Msl3. These evolutionary differences reflect the complex nature of meiosis and highlight the need for species-specific investigations to fully understand the intricacies of this fundamental biological process.

## Acknowledgments

We thank Alicia Mc Carthy for her insightful comments and suggestions. We thank Rico Amato for helping us upload the bioinformatic data in GEO. Dr. E.Z.M Taroc has kindly taken care of the Msl3 mouse line.

## Data availability

RNA seq data are available at GEO (GSE235777). The Msl3flox AKA Msl3^em1Forni^ mouse line is being donated to The Jackson Laboratory.

